# Identification of significant genome-wide associations and QTL underlying variation in seed protein composition in pea (*Pisum sativum* L.)

**DOI:** 10.1101/2024.07.04.602075

**Authors:** Ahmed O. Warsame, Janneke Balk, Claire Domoney

**Affiliations:** John Innes Centre, Norwich Research Park, Colney Lane, Norwich NR4 7UH, UK

**Keywords:** Genome-wide association studies (GWAS), quantitative trait loci (QTL), pea (*Pisum sativum*), seed protein composition, vicilin, legumin

## Abstract

Pea seeds are a valuable source of plant proteins for human and animal nutrition and have various industrial applications. The relative abundance of different seed storage proteins affects protein quality, including digestibility and functional properties of protein extracts. Thus, understanding the genetic basis of seed protein composition is crucial to enhance protein quality and nutritional value through breeding. In this study, we employed two complementary approaches, Genome-Wide Association Study (GWAS) and Quantitative Trait Locus (QTL) mapping, to identify genetic loci underlying seed protein composition in pea. Sodium dodecyl sulfate-polyacrylamide gel electrophoresis (SDS-PAGE) was used to separate the seed proteins, and their relative abundance was quantified using densitometric analysis. For GWAS, we analysed a diverse panel of 209 accessions genotyped with an 84,691 SNP array and identified genetic loci significantly associated with globulins, such as convicilin, vicilin, legumins; and non-globulins, including lipoxygenase, late embryogenesis abundant protein (LEA), and annexin-like protein. Additionally, using QTL mapping with 96 recombinant inbred lines (RILs), we mapped 11 QTL, including five that overlapped with regions identified by GWAS for the same proteins. Within these regions, we identified structural genes for seed proteins and other genes with predicted functions in protein biosynthesis, trafficking, and modification. This comprehensive genetic mapping study serves as a foundation for future breeding efforts to improve protein quality in pea and other legumes.

## INTRODUCTION

Pea seeds contain 20–28% protein ^[1]^ and are increasingly utilised in various plant-based food applications, such as meat, bakery, and alternative dairy products ^[2,3]^. Pea proteins can also be used as functional materials to form biofilms and as coating agents ^[4]^. Unlike soybean and wheat, concerns over allergenic responses are minimal in pea, making it a good candidate crop to support the shift towards plant-based diets and sustainable food production systems. However, plant proteins generally differ from animal proteins in their functional properties, such as amino acid composition and different protein structural classes ^[5]^; therefore, various chemical, enzymatic, and physical modifications are deployed to enhance their behaviour within certain industrial processes ^[6]^. A more sustainable and cost-effective strategy could be to enhance protein composition by breeding and to develop cultivars with protein properties that meet the specific requirements of end users.

Plant seeds contain a mixture of proteins with various physicochemical properties, including overall structure (e.g. globular or non-globular), amino acid composition, and hydrophobicity/hydrophilicity, which in turn influence their functional properties, such as water solubility, gelling properties, water- and oil-holding capacity, and emulsifying and foaming properties ^[6]^. In pea, globulins, a class of globular storage proteins including vicilin, convicilin, and legumin, account for up to 80% of the total seed proteins ^[7,8]^. Additionally, pea seed proteins contain relatively abundant lipoxygenases, albumins, lectins, and trypsin inhibitors ^[1]^. Some of these proteins have been linked to specific nutritional and functional properties. For instance, while legumin contains more sulfur-containing amino acids, which are crucial for human and animal nutrition, vicilin-type proteins have been reported to exhibit better solubility and emulsifying capacity ^[7,9,10]^, as well as higher gelling ability ^[10,11]^. The relationship between protein composition and the quality of the final food products has also been highlighted in soybeans. Poysa et al. ^[12]^ used mutant lines impacted in the subunit composition of glycinin and β-conglycinin, the equivalents of legumin and vicilin in pea, respectively, and they showed that mutating specific protein subunits had significant effects on tofu texture (compression hardness and firmness).

To date, efforts towards genetic improvements in pea protein composition have allowed for enhanced protein digestibility by combining null mutants of lectin A (lecA), pea albumin 2 (PA2), and trypsin inhibitors (TI) ^[13]^. Similarly, CRISPR/Cas9-mediated knockout of the lipoxygenase gene (LOX-2) led to improved flavour profiles in pea seed products ^[14]^. Once other nutritionally and functionally important seed proteins have been defined for both end use and genomically, desirable traits can be stacked into a single genetic background through marker-assisted selection breeding.

Pea seed proteins are genetically complex, with multiple genes encoding the most abundant protein classes. For example, in the Caméor reference genome, 12, 9, 2, 8, and 9 genes encode legumin, vicilin, convicilin, albumin 1 (PA1), and albumin 2 (PA2), respectively ^[15]^. In addition, wide genetic variation in the relative abundance of globulins has been reported among diverse pea accessions ^[8]^. Although the genetic basis of such natural variation has not been well studied, it is likely due to differential gene regulation and the influence of developmental and environmental factors. For instance, in *Arabidopsis*, eight seed-specific transcription factors (TFs) that regulate seed storage protein accumulation have been reported, including four’ master regulators’: LEC1, LEC2, FUS3, and ABI3 ^[16]^. The same TF families have also been identified in chickpea ^[17]^ and wheat ^[18]^ using gene co-expression analyses. In pea, the transcription factor ABA-insensitive 5 (ABI5) has been functionally validated as a key regulator of the relative abundance of a major vicilin ^[19]^. Moreover, environmental conditions such as water stress and nutrient deficiency have been shown to modulate the accumulation of different globulins in pea ^[20]^. Lastly, genetic disruption of other plant processes can have pleiotropic effects on the seed protein composition. For example, mutation of *Starch Branching Enzyme 1* (*SBEI*), known as the *R* locus, leads to reduced legumin and higher protein concentrations, along with a wrinkled seed phenotype, reduced starch, and higher amylose concentrations ^[21–23]^.

Previous studies on pea protein composition have been limited to selected globulin proteins and a relatively small number of pea accessions ^[7–9,24]^. Likewise, genetic mapping of the relative abundance of pea seed proteins is limited to a single study by Bourgeois et al. ^[25]^, who used recombinant inbred lines (RILs) to identify quantitative trait loci (QTL) linked to the abundance of globulin seed storage proteins. In this study, we expanded seed protein composition analyses to 25 abundant seed proteins. We performed genome-wide association studies (GWAS) on a diversity panel of 209 pea accessions and QTL mapping on RILs using high-density SNP markers and identified several genetic loci controlling the relative abundance of pea seed proteins.

## RESULTS

### Protein composition analysis

Understanding the types and proportions of different protein classes in pea seeds is crucial for improving the nutritional value and functional properties of their proteins. For protein composition analysis, SDS-PAGE-based densitometric quantification is considered a high-throughput and cost-effective method and is widely used for large-scale germplasm screening of legumes ^[8,19,20,26,27]^. In this study, we used one-dimensional SDS-PAGE to assess the variation in seed protein composition in a diverse panel of 209 pea accessions, representing a larger global collection of 3140 accessions held at the John Innes Centre. The panel was grown in the field for two seasons and included cultivars, landraces, research lines, and both round- and wrinkled-seeded types. Additionally, 96 recombinant inbred lines (RILs) from two round-seeded parental lines with distinct protein profiles, Caméor and JI0281, were analysed.

We used buffer containing 50 mM Tris and 0.5 M NaCl for total seed protein extraction because seed-abundant globulins are salt-soluble, according to Osborne’s classification ^[28]^. Other seed proteins were also soluble in this buffer, as reflected by the different proteins identified using LC-MS analysis. As for protein identification and quantification, we initially used seven accessions across the diversity clusters described by Jing et al. ^[29]^ to optimise the electrophoresis conditions and identify relatively abundant proteins. From the protein profiles of these lines, 25 consensus protein bands were selected for nano-LC-MS/MS identification. Of these, 14 were globulin storage proteins, with five, four, and five bands belonging to legumin, vicilin, and convicilin, respectively (Figure 1). In addition to lipoxygenases (lipox) and albumin (alb), we identified other non-globulin proteins such as late embryogenesis abundant protein (LEA), putative sucrose-binding protein (SBP), annexin-like proteins, and enoyl-(acyl carrier protein) reductases (ENR1 and 2).

**Figure 1.**
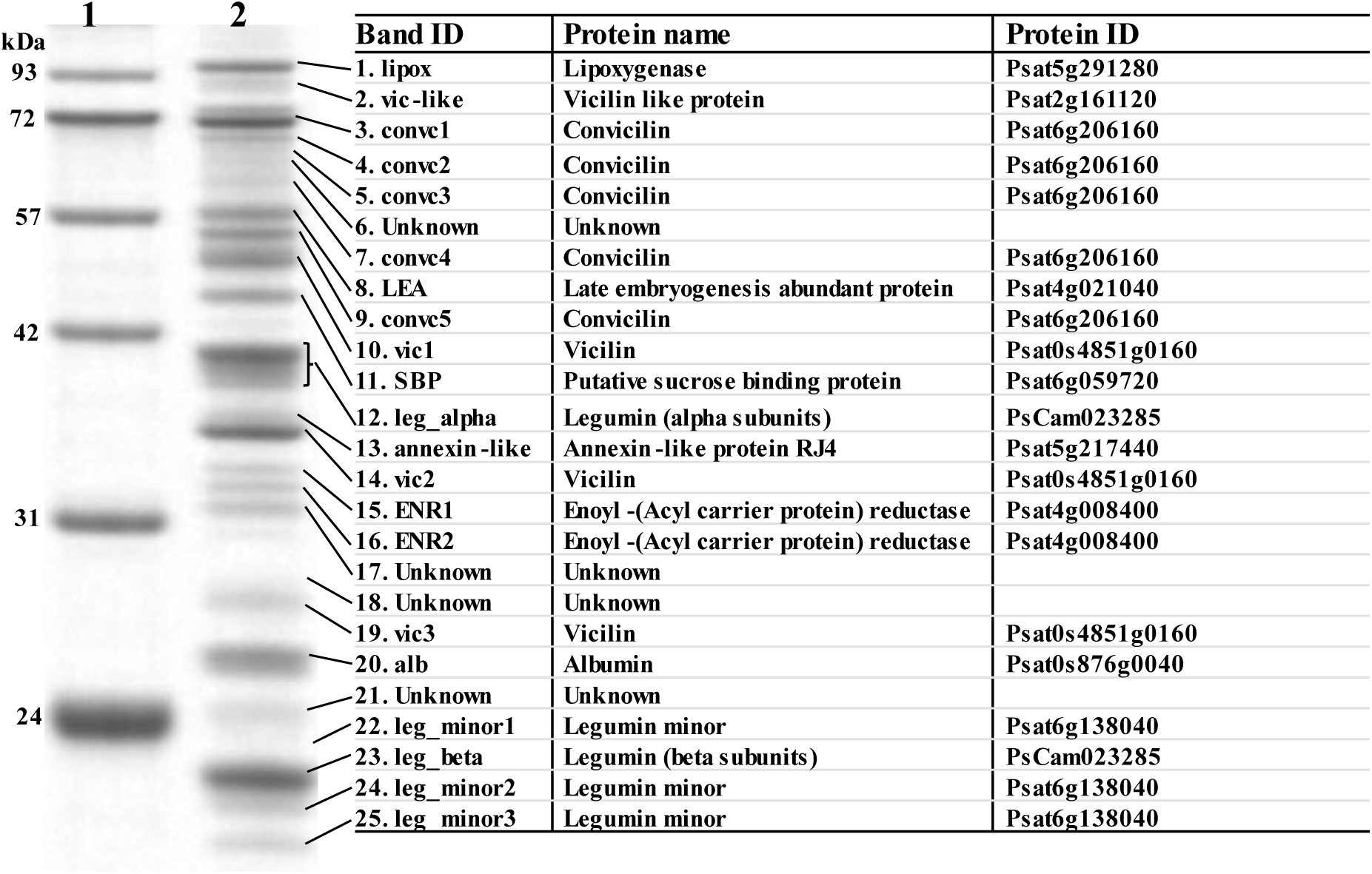
One-dimensional SDS-PAGE profile of pea seed proteins (∼20-100 kDa) run under reducing conditions using 10% Bis-Tris gels. Lane 1: molecular weight standard (Abcam AB116029); lane 2: total seed protein extract from accession number JI0072. Protein identity was determined using non-LC-MS/MS and the Cameor genome v1 (https://urgi.versailles.inra.fr/Species/Pisum/Pea-Genome-project). The short name of the proteins (band ID) is used in this study for convenience, and proteins belonging to the same class are suffixed with numbers based on the order in which they appear on the gel.

Finally, because of the well-known impact of pH on the solubility of seed proteins ^[30]^, we investigated whether changing the pH of the extraction buffer significantly affected the relative abundance of different seed proteins. For this purpose, pea proteins were extracted using eight buffer pH conditions ranging from 3 to 13. Interestingly, the relative abundance of major globulin proteins was largely unchanged across a broad pH range compared to some of the less abundant proteins, such as vic_like, convc3, convc4, leg_minor1, and two unidentified proteins (17_ unknown and 21-unknown) (Figure S1). Overall, these findings suggest that pea proteins are soluble over a broad pH range and that our analysis captures the protein constituents expected in typical pea seeds.

### Variation in protein composition and the impact of *SBE1* mutation

The relative abundance of seed proteins showed considerable variation among the different pea accessions, with a range of more than 4-fold observed for some proteins, including lipox, LEA, convc5, ENR2, and vic3 (Figure 2A). Overall, the most abundant protein bands belonged to legumin, vicilin, and convicilin, which, when individual bands for each class are combined, accounted for 42%, 24%, and 14% of the quantified proteins, respectively. Although the average globulin content found in this study (∼80%) was consistent with that reported in the literature ^[7,31–33]^, the proportion of vicilin was lower than that previously reported for pea ^[7,8]^. This discrepancy is likely due to the prolonged gel electrophoresis step, which was necessary for accurate quantification of the individual protein bands but led to the exclusion of low-molecular-weight peptides (<20 kDa), including several post-translationally processed vicilin peptides from the gels.

**Figure 2.**
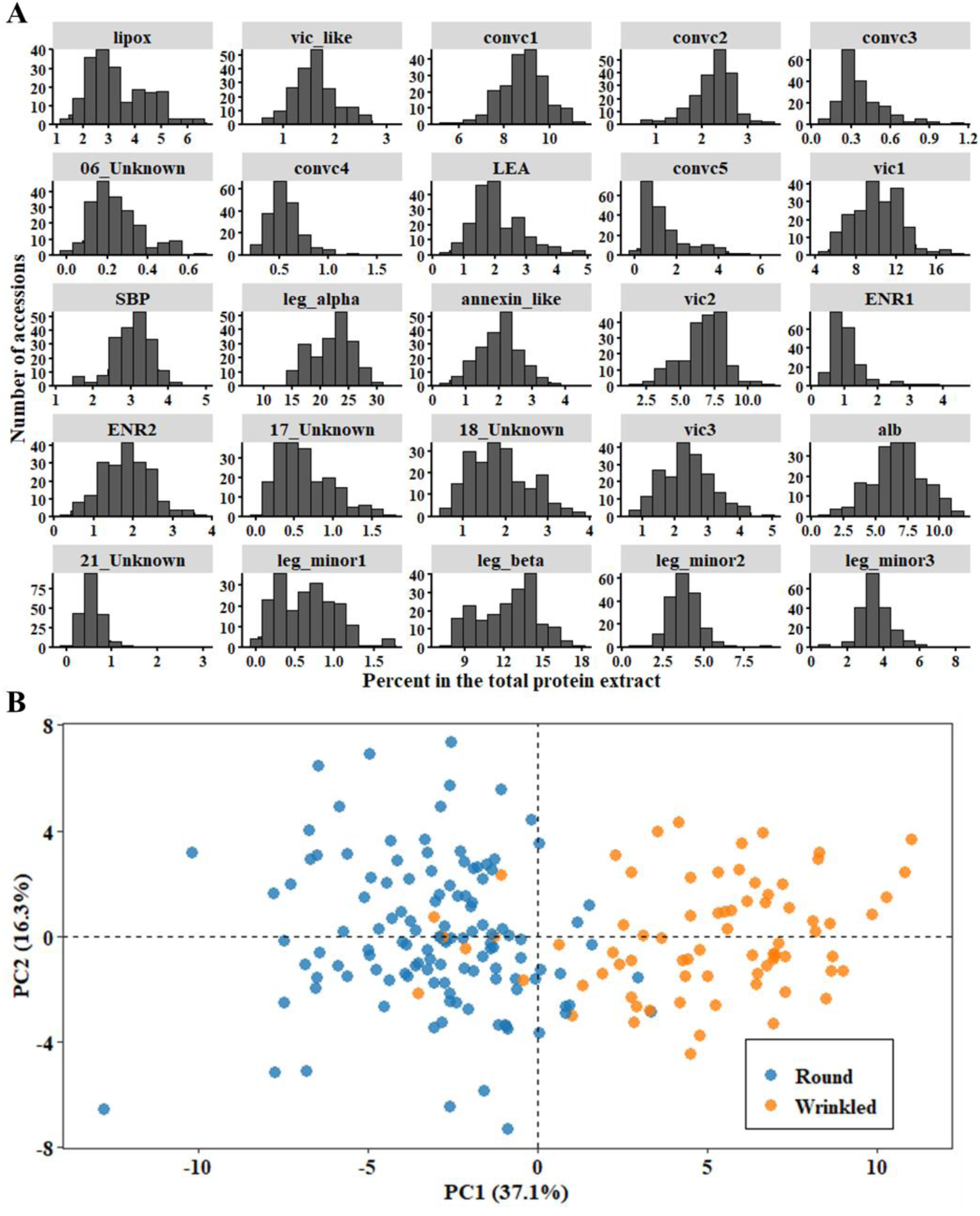
Variation in seed protein composition in the pea diversity panel. (A) Histograms of the relative abundances of 25 polypeptides. The data were based on the adjusted means of 196 accessions grown in the field during 2021 and 2022. Relative proportions were calculated densitometrically for each protein band based on the total protein intensity in each gel lane. (B) Principal component analysis based on the data in (A), showing two clusters representing round and wrinkled (*sbe1* or *rr*) accessions.

Of the 209 accessions in the diversity panel, 77 were wrinkled accessions, which is a common trait in cultivated vining (vegetable) peas. Based on the results of a previous screen of the JIC pea collection by Rayner et al. ^[34]^, all wrinkled accessions carrying the so called *r* allele of the *R* locus are homozygous for a well-characterised mutation in *Starch Branching Enzyme 1* (*SBEI*) ^[23]^. The relative abundance of the main protein bands differed markedly between the round and wrinkled peas (Figure 2B). Our results show that the reported reduction in the major legumin polypeptides in wrinkled peas ^[21,22]^ is compensated by the over-accumulation of less abundant proteins, including lipoxygenase (lipox), LEA, convc5, SBP, ENR, and albumin (alb, also known as PA2). These relationships were further explored using pairwise correlation analysis (Figure S2), in which the aforementioned proteins showed a significant negative correlation with another group of proteins, including leg_alph, leg_beta, vic3, and leg_minor1. Conversely, the relative abundance of convc 1, 2, and 4 and annexin-like proteins was not affected by the *SBEⅠ* mutation.

To further investigate whether the differences in protein composition correlated with significant changes in amino acid profiles, we selected four round and four wrinkled pea accessions for amino acid analysis. In addition to the seed phenotype, these accessions differed in the relative abundance of specific proteins (Figure S3A). As expected, there were small but statistically significant differences (*p* ≤ .05) among accessions in 10 of the 16 amino acids quantified in this study. However, the most prominent difference was between the two seed groups, where wrinkled accessions had significantly higher levels of methionine and lysine (Figure S3B).

### Genome-wide association analysis of protein abundance

To determine the genetic basis of the variation in the relative abundance of soluble seed proteins, we performed genome-wide association analyses between the quantified band intensity of of individual proteins and 32,972 high-quality SNPs. To reduce false positives due to population stratification, the population structure was inferred using fastSTRUCTURE ^[35]^ and the LEA R-package ^[36]^. Both software programs predicted 18 subpopulations (Figure 3A, B), and we included them as covariates in the GWAS analysis. The complex admixture in the population was consistent with the selection of these accessions across 17 subclusters within the three major diversity groups reported in the John Innes Centre pea collection ^[29]^. In contrast, the first two principal components (PCs) accounted for only 68.5 % of the genetic variation and showed a less clustered structure compared to fastSTRUCTURE (Figure 3C). The ancestry data from fastSTRUCTURE software were used as covariates in the GWAS analysis.

**Figure 3.**
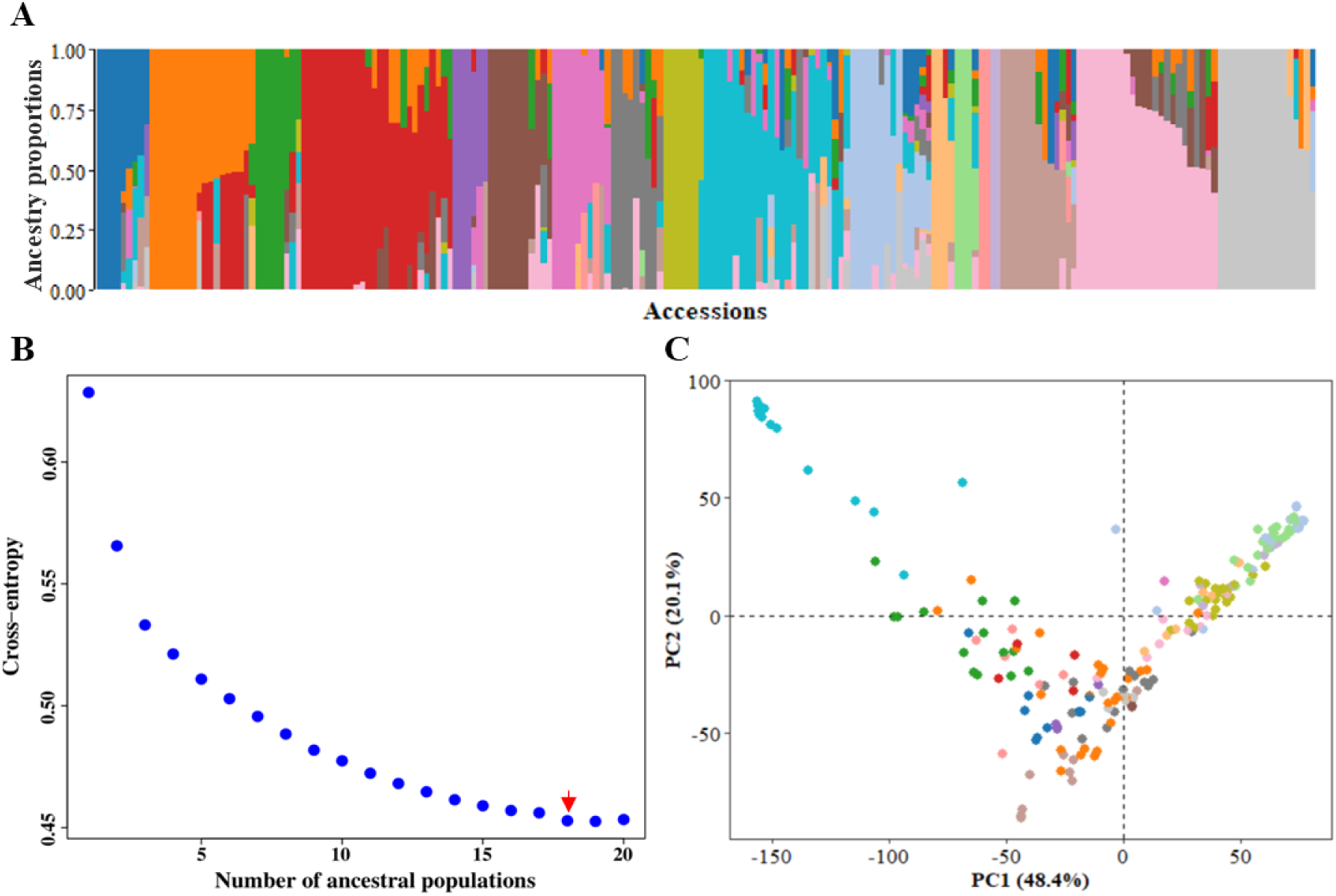
Population structure of the 209 pea accessions used in this study. (A) Ancestry proportions predicted using fastSTRUCTURE software. (B) Population genetic structure predicted by the snmf algorithm in the LEA R package, with k ranging from 1 to 20. The k value with the lowest cross-entropy (k = 18, marked by a red arrow) represents the number of sub-populations in the panel. (C) Population stratification based on principal component analysis, where accessions are coloured according to clusters identified by fastSTRUCTURE software.

Several GWAS analysis methods with varying computational efficiency, statistical power, and control of false positives have been developed over the years, and commonly used association analysis software, such as GAPIT ^[37]^, TASSEL ^[38]^, and rMVP ^[39]^, allow for the implementation of multiple GWAS models. The main advantage of combining different models is ensuring that important loci are not missed because of the power limitations of an individual GWAS model. Therefore, we performed association analysis using BLINK, FarmCPU, and SUPER. As shown by the quantile-quantile plots (see Figures 5, 6, and 7), the BLINK model showed the least deviations from the expected p-values, whereas FarmCPU showed suppressed p-values to control for false positives. In contrast, SUPER exhibited inflated p-values for most of the traits. In a recent comparison of commonly used GWAS models, Cebeci et al. ^[40]^ found that BLINK and FarmCPU were the best-performing models for controlling false positives and identifying QTL, with BLINK being more powerful in detecting those with lower heritability. The combined list of significant SNP associations found across the two years for 24 polypeptides is given in Table S2.

### Association of protein composition with the *R* locus

Consistent with the difference in protein profiles of round and wrinkled pea seeds (Figure 2B), 51 SNPs in the 90-120 Mb region of chromosome 3) harbouring the mutation in *SBE1* (*R* locus) were associated with 14 out of 25 polypeptides, as detected by at least one of the GWAS models in either year (Figure 3A). The most significant association in both years was recorded for the abundance of convicilin (convic5) with SNP AX-183861246 located at 95,634,854 bp. Other proteins highly associated with SNPs in this region included vicilin (vic3), legumin (leg_alpha), lipoxygenase (lipox), and late embryogenesis abundant protein (LEA). Surprisingly, these significant SNP associations were spread across a chromosomal segment spanning over 30 Mb, with the most significant SNPs being nearly 10 Mb from the physical location of the *R* locus (Figure 3A). A similar broad GWAS peak for the *R* locus has recently been identified in this region using a much larger diversity panel ^[41]^.

To investigate this region further, we compared linkage disequilibrium (LD) between the SNPs within 95–120 Mb of chromosome 3 to further investigate this region using a subset of 72 accessions each from the wrinkled-and round-seeded pea groups. Unlike round pea, wrinkled accessions showed high correlations between SNPs and long-range LD blocks (compare Figure 4B (wrinkled) with Figure 4C (round)), which may reflect a high selection pressure for other traits in this region. Wrinkled peas are generally sweeter because of the higher sucrose content associated with the *sbeI* allele ^[23]^; therefore, they are primarily bred for frozen pea market together with other desirable traits. It is worth noting that this region contains two other agronomically important loci: *Det,* which controls indeterminacy ^[42]^, and *Tl,* which controls the formation of tendrils ^[43]^. The selection for these morphological traits has likely contributed to the difference in LD blocks between the round and wrinkled accessions. Furthermore, the gene encoding the transcription factor ABA-insensitive 5 (*ABI5*), which plays a central role in the regulation of vicilin accumulation in *Medicago* and pea ^[19]^, is also located in this region (Figure 4A).

**Figure 4.**
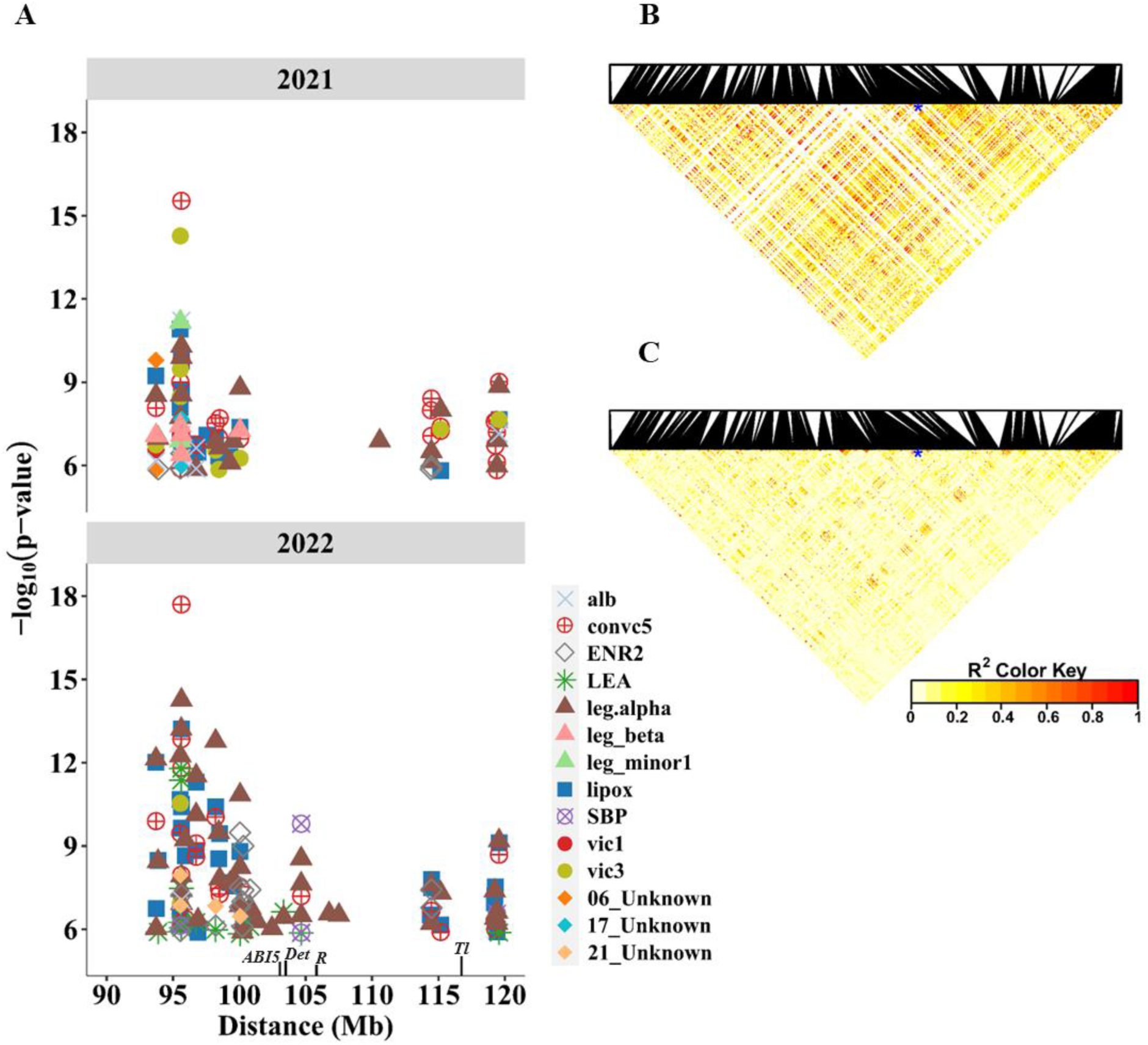
SNPs associated with 14 abundant seed proteins are located on chromosome 3 flanking the *R* locus. (A) The 90 – 120 Mb segment of chromosome 3 showing the positions of SNPs significantly associated with different proteins during 2021 and 2022. Also shown is the location of *ABI5*, *Det*, *R*, and *Tl*, which control globulin composition, indeterminacy, seed shape, and tendrils in pea, respectively. (B, C) Heatmaps showing the linkage disequilibrium (LD) of SNPs within the chromosomal segment based on 72 accessions of wrinkled and round pea. LD is expressed as the squared correlation (R^2^) between the alleles of SNPs, and the red colour in the heatmap denotes regions with a high LD. The blue asterisks in the heatmaps indicate the location of the SNP closest to the *R* locus.

### Loci associated with globulin proteins

Convicilin is one of the predominant globulin proteins in pea seeds, accounting for approximately 14% of the quantified proteins, with five polypeptides of different molecular weights (∼51-70 kDa). This protein is closely related to vicilin, except for the presence of an N-terminal extension, which is rich in hydrophilic and highly charged residues ^[44,45]^, and the presence of methionine and cysteine residues ^[46]^. Genome-wide association analysis identified a region at the end of chromosome 5 (∼593-600 Mb) that was significantly associated with three of the convicilin polypeptides (convc1, convc2, and convc3) during 2021 (Figure 5, Table S2). Interestingly, this region contains an orthologue of a C3H family transcription factor (Psat5g273280,∼ 597.5 Mb), which showed high co-expression (r = 0.98) with seed storage proteins in chickpea ^[17]^. In addition, in the 2022 season, convc3 and convc5 were associated with SNPs near convicilin structural genes Psat6g206160 and Psat6g206200 on chromosome 6 (∼458.3 - 458.4 Mb). On the other hand, different from other convicilins, the second most abundant convicilin, convc5, was highly associated with SNPs at the *R* locus on chromosome 3 (Figure 5C), suggesting that this convicilin isoform is distinct in its genetic control, and matches the previously described A-type convicilin with a molecular mass of ∼ 60 kDa ^[44]^.

**Figure 5.**
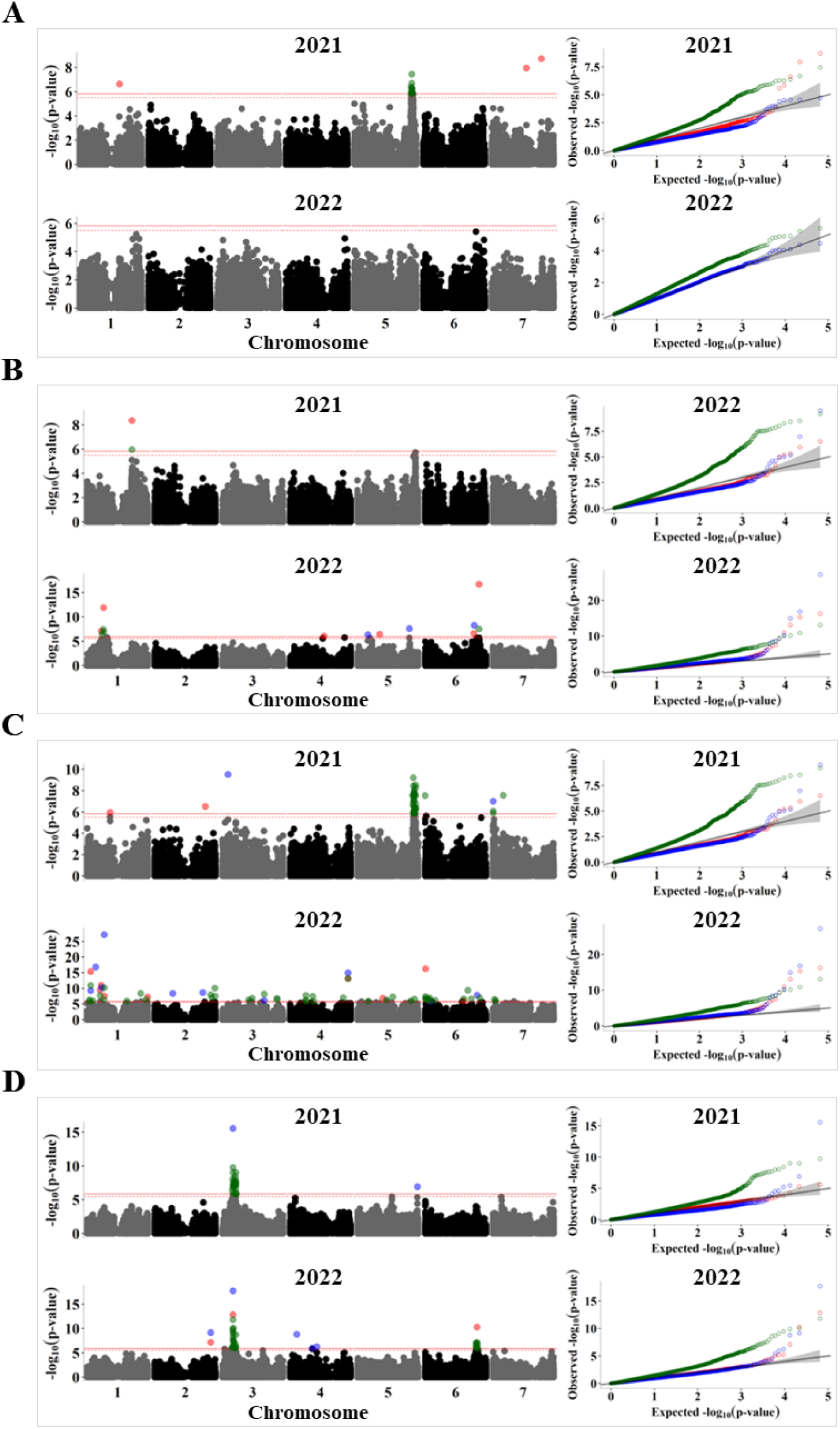
GWAS analysis of the relative abundance of convicilin polypeptides during 2021 and 2022. Manhattan plots (left) and quantile-quantile (QQ) plots (right) for (A) convc1, (B) convc2, (C) convc3, and (D) convc5. The solid horizontal line is the GWAS significance threshold corresponding to the false discovery rate (FDR) at p = 0.05, whereas the dotted line is the threshold for suggestive association at p = 0.1. SNPs above the significance threshold in the Manhattan plots and the QQ plots are coloured by GWAS model, with red, blue, and green denoting results from BLINK, FarmCPU, and SUPER, respectively.

Vicilin is the most abundant globulin in pea seeds, and we quantified three gel bands with vic1 (∼48 kDa) being the most abundant polypeptide. In addition to the significant GWAS signals identified in 2022 on chromosome 5 near the loci associated with convicilins (Figure 6A), five SNPs at 512-516 Mb were detected on chromosome 6 in both years. However, the most consistent peak identified in both seasons by the SUPER GWAS model was a region on chromosome 3 (250-252 Mb), linked to vic2 (Figure 6B). This region contains the annotated vicilin structural genes Psat0s4055g008 (not assigned to a chromosome in the Caméor reference) and Psat3g104920. The low-molecular-weight vicilin polypeptides vic2 and vic3 of ∼36 and ∼27 kDa, respectively, are predicted to result from post-translational cleavage of the primary translational products ^[47]^; thus, genes related to protein processing can be potential candidates for variation in their relative abundances. Similar to convc5, vic3 was primarily associated with SNPs at the R locus on chromosome 3 (Figure 6C), but they were affected differently. While convc5 was over-accumulated in wrinkled pea accessions, vic3 was more abundant in round peas.

**Figure 6.**
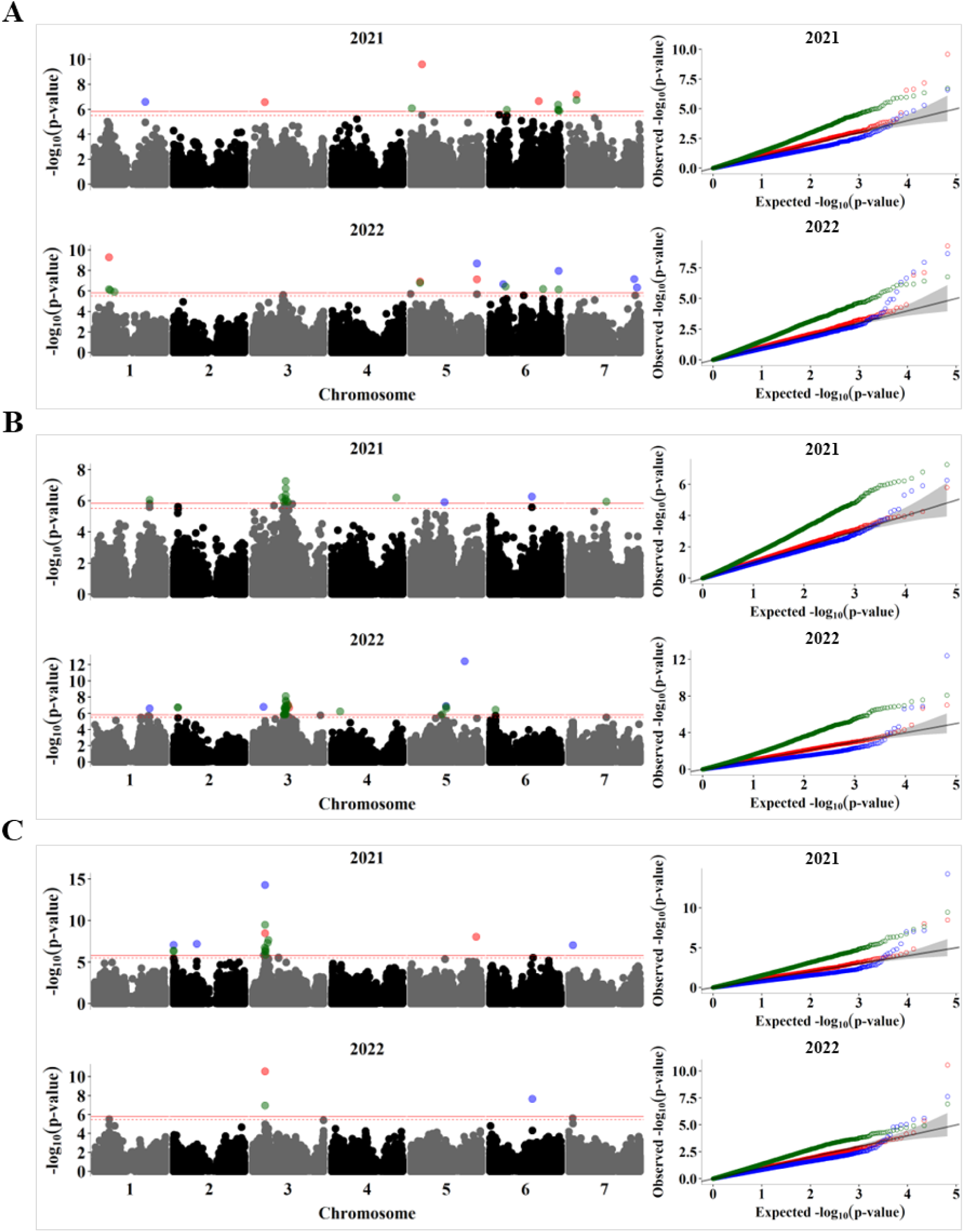
GWAS analysis of the relative abundance of vicilin polypeptides in 2021 and 2022 Manhattan plots (left) and quantile-quantile (QQ) plots (right) for (A) vic1, (B) vic2, and (C) vic3. The solid horizontal line is the GWAS significance threshold corresponding to the false discovery rate (FDR) at p = 0.05, whereas the dotted line is the threshold for suggestive association at p = 0.1. SNPs above the significance threshold in the Manhattan plots and the QQ plots are coloured by GWAS model, with red, blue, and green denoting results from BLINK, FarmCPU, and SUPER, respectively.

For legumins, although their abundance was primarily associated with the *R* locus on chromosome 3 (Figure S4, Table S2), significant SNP associations with leg_alpha, leg_beta, and leg_minor1 were mapped across all chromosomes. However, there was no overlap between the regions identified during these two seasons. Considering the importance of legumins as major seed storage proteins and the confounding effect of the *R* locus, we performed an association analysis for a subset of 130 round-seeded accessions. We did not find any significant associations for legumin proteins, likely owing to the loss of statistical power because of the smaller population size and the multigenic nature of this class of proteins.

### Loci underlying non-globulin pea proteins

Non-globulin proteins are relatively abundant in pea seeds and influence their nutritional quality ^[13,14]^. Albumin is the most abundant non-globulin protein in pea seeds, and is composed of albumin 1 (PA1) and the higher molecular weight albumin 2 (PA2) shown in Figure 1 at ∼26 kDa and is referred to as alb in this paper. Although the abundance of this protein (PA2) was primarily associated with the *R* locus, two regions on chromosomes 1 (350.3-350.8 Mb) and 4 (58.9-62.4 Mb) were identified in 2022 (Figure 7A).

**Figure 7.**
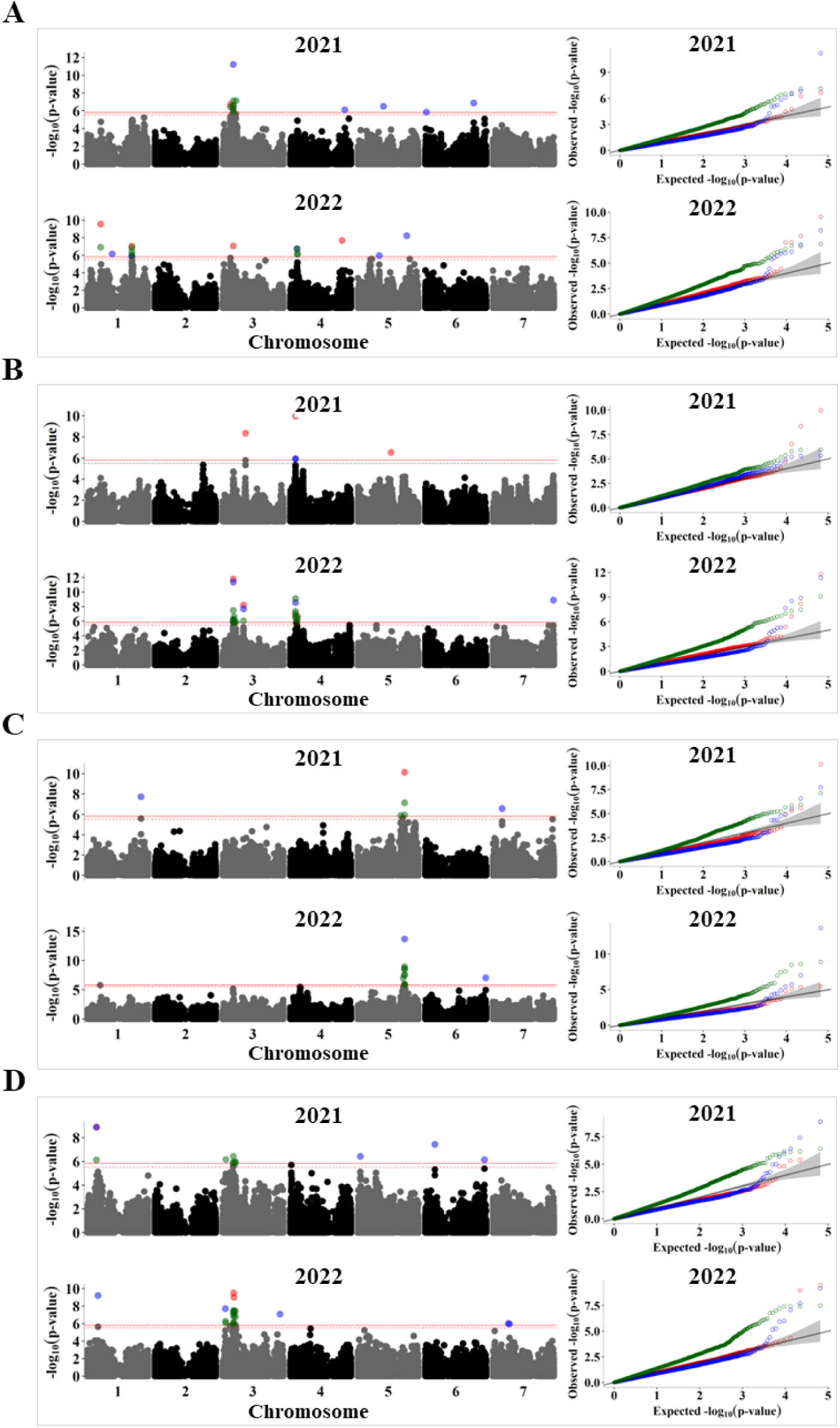
GWAS analysis of relative abundance of non-globulin seed proteins during the 2021 and 2022 seasons. Manhattan plots (left) and quantile–quantile (QQ) plots (right) for (A) alb, (B) LEA, (C) annexin-like, and (D) ENR2. The solid horizontal line is the GWAS significance threshold corresponding to the false discovery rate (FDR) at p = 0.05, whereas the dotted line is the threshold for suggestive association at p = 0.1. SNPs above the significance threshold in the Manhattan plots and the QQ plots are coloured by GWAS model, with red, blue, and green denoting results from BLINK, FarmCPU and SUPER, respectively

Late embryogenesis abundant proteins (LEA) and annexin-like proteins are among the least studied proteins in the context of seed protein composition but are relatively abundant in the seed. Both proteins belong to multigene families and are required for environmental adaptation in plants ^[48,49]^. We identified a region on chromosome 4 (∼44.9-45.9 Mb), which was significantly associated with the abundance of LEA protein in both years (Figure 7B). A significant SNPs on the same chromosome, at 58.7 Mb, colocated with a structural gene for LEA Psat4g026680. As for annexin-like protein, significant SNP associations were located on chromosome 5 at ∼496-498.6 Mb in both years. This locus harbours two structural genes annotated as annexin-like proteins (Psat5g217920 and Psat5g217440 at 498.64 and 498.8 Mb, respectively).

Among the metabolic changes in pea seeds associated with the *R* locus, increased lipid content has been observed in wrinkled pea ^[23,50]^. We found that polypeptides belonging to the enoyl acyl carrier protein reductase enzyme (ENR), the last enzyme in the fatty acid elongation cycle, were more abundant in wrinkled pea accessions. GWAS analysis revealed an association peak for ENR1 and ENR2 on chromosome 1 (71.3-72.7 Mb), which colocates two plant lipid transfer genes, Psat1g038160 and Psat1g038200. Another GWAS signal was detected in both years on chromosome 3 (26.8-29.7 Mb) and was associated with ENR2. Other proteins with significant associations included a putative sucrose-binding protein (SBP), a vic-like protein, and four unidentified seed proteins (Table S2).

### QTL mapping of abundant seed proteins in round-seeded RILs

To validate the significant GWAS signals and discover new loci underlying variations in protein composition, we used 96 recombinant inbred lines from a cross between Caméor and JI0281. JI0281 is an Ethiopian landrace with a very different protein profile from Caméor (Figure S6). Both Caméor and JI0281 are round-seeded; therefore, the *R* locus had no confounding effect on the protein composition. Using composite interval mapping in the R/qtl package with 17k SNP markers distributed on seven linkage groups (LG) representing the seven pea chromosomes, 11 QTL linked to 14 protein bands were mapped on five different LGs (Figure 8). By mapping the LGs to their chromosome-level genome assembly, we found that five QTL overlapped regions containing significant GWAS associations for the same proteins (Table 1). On linkage group 6 (1LG6), corresponding to chromosome 1, a QTL at 33.3-34.3 centimorgan (cM) was strongly linked to ENR2 and other unidentified protein (17_Unknown). This region also contained a significant SNP identified by GWAS, which explained 21% of the phenotypic variance in ENR2 in 2021. Another QTL for this protein on 6LG2 colocated with QTL for a putative sucrose binding protein (SBP) and an unidentified protein band (18_Unknown). On the other hand, we mapped QTL on 6LG2 (160.36-162.85 cM), 5LG3 (170.83-171.54 cM), and 3LG5 (104.03-105.89 cM) that were linked to convicilin (convc5), annexin-like, and vicilin (vic2), and explained 30, 27 and 47% of the phenotypic variation, respectively (Figure 8, Table 1). These QTL intervals overlapped regions containing strong GWAS signals and known structural genes for these proteins. For vicilin-like, a QTL interval 96.95-100.48 on 1LG1 (∼ 458.6-465 Mb on chromosome 1) was linked to the coding gene at ∼ 465.5 Mb. Finally, individual QTL were identified for lipoxygenase (lipox) and convicilin (convc2) (Figure 8), whereas no QTL were detected for the relative abundance of any legumin polypeptide.

**Figure 8.**
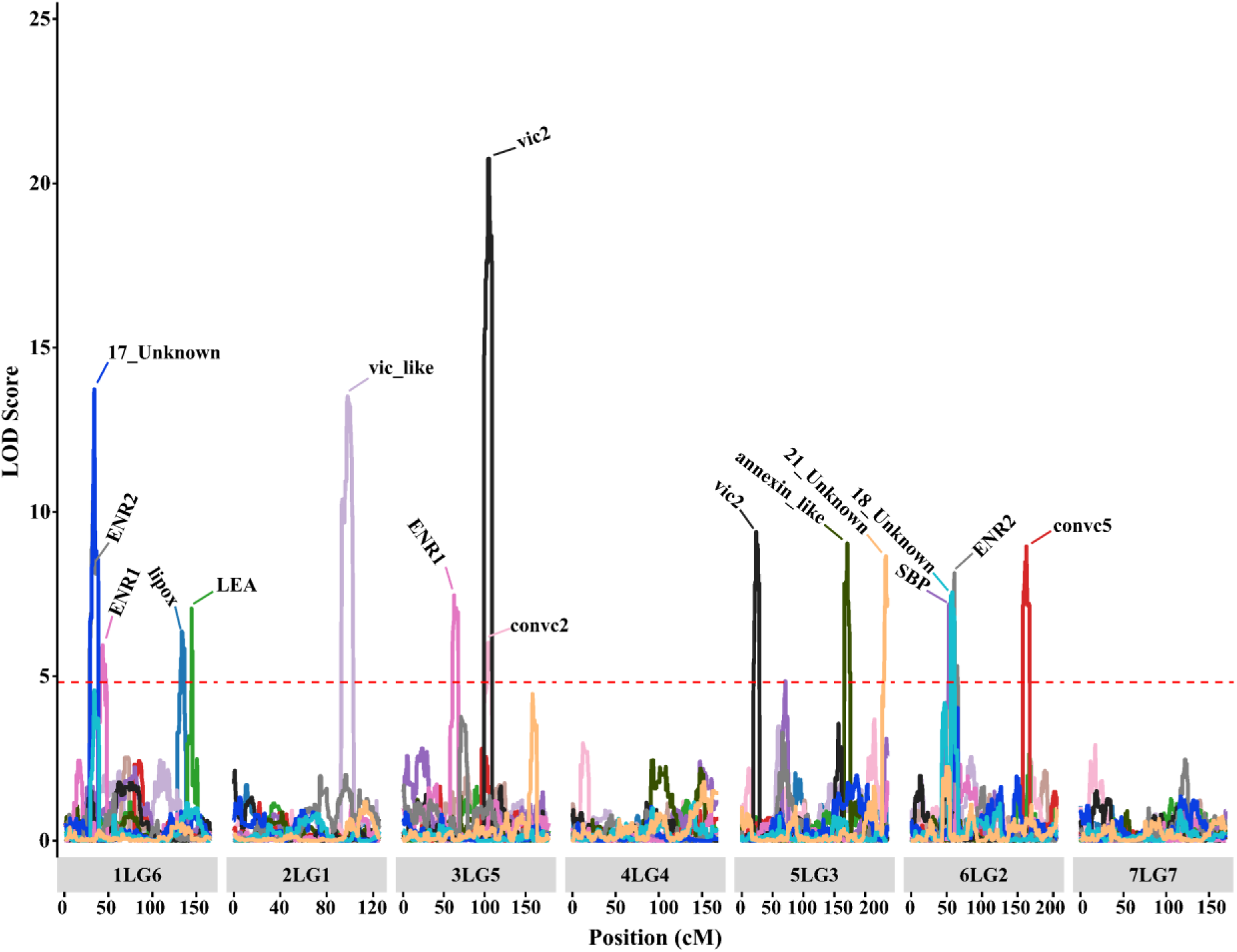
QTL identified for protein composition in a JI0281 × Caméor RIL population of 96 lines. Chromosome numbers and linkage groups are shown on the x-axis for ease of comparison with other studies (for example, 1LG6 indicates chromosome 1 and the corresponding linkage group 6 in the genetic map). The horizontal dashed line indicates the LOD threshold for significance at alpha ≤ 0.05. See Table 1 for details on the QTL mapping summary.

**Table 1.**
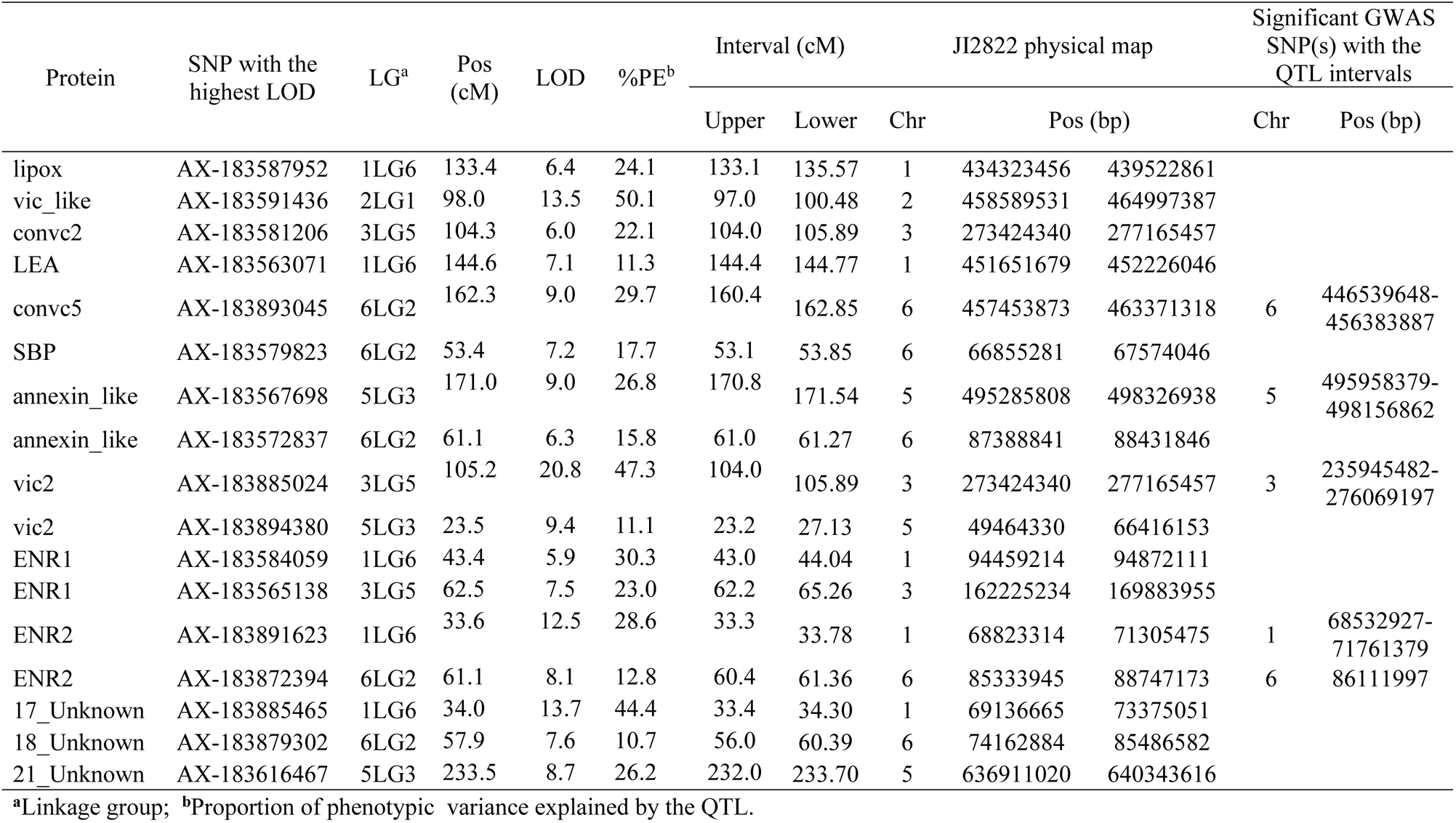
Positions of QTL (quantitative trait loci) linked to relative abundance of different seed proteins in pea using 96 RILs from Cameor × JI281.

## DISCUSSION

Pea is a key source of plant-based proteins for human and animal nutrition, particularly in temperate climates unsuitable for soybean or *Phaseolus* cultivation, such as Northern Europe and parts of Canada, where this crop is well adapted. From a nutritional and quality perspective, a key objective in pea breeding, driven by recent market growth in plant-based proteins, is to improve protein composition to meet the quality requirements of various end-uses. The coding genes for several major pea seed proteins are known ^[1,51–53]^, and some of their physicochemical properties have been documented ^[7,10,11,45]^. This study describes the natural diversity and reveals the genetic determinants of the relative abundance of pea seed proteins using combined GWAS and QTL analyses.

Protein composition analysis revealed variations in the abundance and subunit composition of different seed proteins. For instance, convicilin has two known coding genes, Psat6g206160 and Psat6g206200 ^[15]^; however, a single protein band of approximately 70 kDa on SDS-PAGE gels is generally referred to as convicilin ^[7,8,10]^. Our analyses using nano-LC-MS identified five polypeptides of convicilins (∼51-70 kDa), with the most abundant being convc1 (∼70 kDa) and convic5 (∼53 kDa), with the exception of the JI0960 line, which had convic3 (∼64 kDa) as the predominant convicilin. In addition, unlike major conv1, the second most abundant band (convic5) was significantly affected by the *R* locus and over-accumulated in the wrinkled peas. The presence of multiple polypeptides and the difference between conv1 and conv5 in relation to the *sbeI* mutation may indicate that convicilins may be more heterogeneous than previously thought, and further investigation is needed to determine the basis of such heterogeneity.

We found a marked difference in protein composition between round and wrinkled peas and pairwise correlations between different proteins, which may provide insight into their intricate relationships and which proteins to target for genetic improvement in each group. For instance, field pea (round-seed) is mainly used in the animal feed industry, where increasing the content of cysteine and methionine can significantly enhance the nutritional value of animal feeds. It is well known that legumin and albumin have a higher content of these amino acids ^[1,20]^, and silencing the accumulation of other seed proteins, such as lipoxygenase, late embryogenesis abundant protein (LEA), SBP, and annexin-like protein, can potentially increase their proportion. In this context, our results provide the critical genetic information required for such an approach in pea. It is worth noting that direct knockout of genes encoding major storage proteins could lead to drastic proteome rebalancing where previously less abundant proteins are overaccumulated, as reported in soybean mutants lacking β-conglycinin and glycinin ^[54]^.

The vast majority of GWAS peaks did not co-localise with structural genes, with the exception of convc5, LEA, annexin-like, and vic2, which is consistent with the previously reported involvement of trans-regulatory loci in seed protein composition ^[19,25]^. We investigated the gene content of the regions flanking significant SNPs that explained more than 10% of the phenotypic variance, and found that several genes could be related to variations in protein composition, including protein biosynthesis, trafficking, and modification (Table S3). However, the role of these genes in the abundance of seed proteins needs to be determined by either co-expression analysis or mutant studies.

The integration of GWAS and QTL results provided further support for the identified genomic regions influencing seed protein composition and validated five loci identified by GWAS. Bourgeois et al. ^[25]^ used two RIL populations involving Caméor and two other French cultivars, VavD265 and Ballet, and identified 365 protein quantity loci (PQLs) for 245 spots on 2D gels. In this study, we used JI0281, which is a small seeded landrace from Ethiopia. Unfortunately, our attempt to overlay the two studies and find consensus QTLs was hindered by the dispersed linkage groups and low marker coverage in Bourgeois et al. ^[25]^.

Overall, our comprehensive analysis offers valuable insights into the genetic architecture of seed protein composition in peas, providing a foundation for future research aimed at dissecting the molecular mechanisms underlying trait variation and facilitating targeted breeding efforts for improved protein quality in pea cultivars. Further functional characterisation of candidate genes within the identified genomic regions will be essential to elucidate their specific roles in modulating seed protein composition and to inform strategies for enhancing nutritional quality and agronomic performance in pea.

## METHODS

### Plant material

The diversity panel used for GWAS analysis, known as the PCGIN panel, comprised 230 pea accessions selected across the diversity spectrum of *Pisum* ^[29]^. The panel was grown in 0.5 m^2^ plots, with 40 plants per plot, for two growing seasons, during early summer 2021 and 2022, at the Dorothea de Winton Field Station in Bawburgh, Norfolk, UK. Crop management practices included insecticide and fungicide sprays and 2 – 3 treatments with 15% manganese during the growing season. Upon maturity, all plants in the plot were harvested, and the seeds were pooled for further analysis. Only accessions with sufficient seeds were used, namely 196 lines from the 2021 harvest and 209 lines from 2022.

For QTL analysis, we used 96 F_7_ recombinant inbred lines (RILs) derived from a wide cross between the French cultivar Caméor and JI0281, an Ethiopian landrace. RILs were grown in a glasshouse during the spring/summer period of 2022, as described by Ellis et al. ^[55]^. Briefly, individual plants were grown in 9 cm diameter pots containing a peat/loam/grit mix (65:25:10) supplemented with 3 kg/m^3^ dolomitic limestone. The plants were supported by canes and watered twice daily using an automated watering system. At maturity, seeds from individual F_7_ plants were harvested, pooled, and analysed for protein composition.

### Protein extraction and SDS-PAGE analysis

Approximately 5 g of dry mature pea seeds was ground using an IKA Tube mill (IKA, Staufen, Germany) and sieved with a 0.25 mm Nylon mesh screen. For total protein extraction, seed flour (approximately 30 mg) was mixed with 1 ml of extraction buffer (50 mM Tris-HCl, pH 8, containing 500 mM NaCl) for 2 h at 4°C. After centrifugation for 10 min at 15700 × *g* at 4°C, the supernatants were collected and diluted with water at a 1:4 ratio, and the protein concentration was determined using a Direct Detect® Infrared Spectrometer (Millipore, Merck).

For SDS-PAGE separation, protein samples were mixed with loading buffer (125 mM Tris-HCl, pH 6.8, 20% SDS (w/v), 4% glycerol (v/v), 10% β-mercaptoethanol (v/v), and 0.5 mg/ml bromophenol blue) and heated at 80 °C for 10 min. Approximately 2.5 µg of protein was loaded into each well of precast 10% NuPAGE Bis-Tris gels (Invitrogen, Thermo Fisher Scientific). Each gel contained two control samples, which were repeated across all gels to capture gel-to-gel variation. In the RIL population, the test lines were replicated twice in separate gels and the parental lines were used as control samples for each gel. After running at 200 V for 75 min using NuPAGE MOPS SDS Running Buffer (Invitrogen, Thermo Fisher Scientific), the gels were stained with 25 ml of InstantBlue Coomassie protein staining solution (Abcam, Cambridge, UK) overnight and rinsed with water for at least 16 h.

### Identification of seed protein bands

A standard in-gel digestion protocol followed by nano-LC-MS/MS analysis was used to identify seed protein bands. Briefly, the slices were washed with 50 mM triethylammonium bicarbonate (TEAB) buffer (pH 8; Sigma-Aldrich, Merck), incubated with 10 mM DTT in 50 mM TEAB for 30 min at 65 °C, and finally with 30 mM iodoacetamide in 50 mM TEAB at room temperature. After washing and dehydration with acetonitrile, gel slices were soaked in 50 mM TEAB containing 10 ng/µL sequencing-grade trypsin (Promega, Southampton, UK) and incubated at 50 °C for 8 h. The extracted peptide solution was dried, dissolved in acetonitrile containing 0.1% trifluoroacetic acid (v/v), and analysed using an Orbitrap Eclipse Tribrid mass spectrometer coupled with an UltiMate^®^ 3000 RSLCnano LC system (Thermo Fisher Scientific, Hemel Hempstead, UK). The acquired MS spectra were searched against a protein sequence database for *Pisum sativum* compiled from the INRA Caméor genome project (https://urgi.versailles.inra.fr/Species/Pisum/Pea-Genome-project), JI2822 genome (in-house database), and the Caméor Unigene database ^[56]^.

### Densitometric quantification of protein bands

Images of the Coomassie Blue-stained gels were captured with a G:box Chemi XRQ (Syngene, part of Synoptics, Cambridge, UK), processed with ImageJ (https://imagej.net) to remove the background using a rolling ball radius of 50 pixels, and cropped to the desired dimensions. The images were then imported into GelAnalyzer software (http://www.gelanalyzer.com/) for densitometric quantification of protein bands. After the software automatically detected the gel lanes and bands, each lane and band was manually checked and adjusted if required. In addition, the background detection parameter was set to 10% using the rolling-ball method. The percentage of each band was calculated as the intensity value of that specific band divided by the total intensity of all bands in each lane, multiplied by 100.

### Data analysis

Due to the different numbers of accessions evaluated in each growing season (196 and 209 accessions in 2021 and 2022, respectively), the data were analysed separately. ANOVA was conducted using R (R Core Team), and genotype means adjusted for gel batch effects were obtained using the emmeans R package to calculate the estimated marginal means ^[57]^. In addition, the data for most protein bands were not normally distributed; therefore, data transformation was performed using inverse-normal transformation according to the formula given by Yang et al. ^[58]^. To calculate the correlation coefficients between the relative abundance of seed proteins, data from 196 lines grown in both seasons were used.

### SNP genotyping

DNA was extracted from young leaves of the diversity panel using an in-house DNA extraction protocol. Briefly, leaf tissue was ground in liquid nitrogen and homogenised with 400 µL of extraction buffer (0.5 M NaCl, 0.1 M Tris-HCl, 0.05 M EDTA, pH 8). Next, 4 µL RNase (QIAGEN, Venlo, The Netherlands) was added, and the mixture was incubated at 65 °C for 15 min. This was followed by the addition of 20 µL of 20% SDS and 400 µL of phenol/chloroform/IAA (25/24/1), and centrifugation at 20,000 × *g*. The clear supernatant was collected and DNA was precipitated with 1 ml of 100% ethanol, followed by two washes with 70% (v/v) ethanol. For the RIL lines, DNA was extracted using the oKtopure kit (Biosearch Technologies, Middlesex, UK), according to the manufacturer’s instructions. Finally, the DNA quantity and quality were checked using a Nanodrop and sent to Neogen, Ltd. (Neogen Europe, Ayr, Scotland) for genotyping using a pea axiom array containing 84,691 SNPs ^[55]^.

### GWAS analysis

Raw genotypic data were subjected to quality control using PLINK software ^[59]^. The missing data threshold was set to 10%, and SNP markers with a minor allele frequency ≤ 5% were removed. This resulted in a final list of 32,829 SNPs anchored on the physical map positions of the cultivar Caméor and the draft genome assembly of JI2822 (Table S1). These data were then used to predict the population structure of the pea diversity panel using fastSTRUCTURE (Raj et al., 2014) and the snmf function in the LEA R package (Frichot & François, 2015). Additionally, principal component analysis was performed using the GAPIT3 R package ^[37]^. Marker-trait association analysis was performed in the GAPIT3 R package ^[37]^ using three models: Bayesian information and linkage disequilibrium iteratively nested keyway (BLINK), fixed and random model circulating probability unification (FarmCPU), and settlement of MLM under progressively exclusive relationship (SUPER). The first two are multi-loci, whereas SUPER is a single-locus model which has improved statistical power compared to similar GWAS models ^[60]^. A Bonferroni-corrected threshold of 5.82 and 5.5 (–log_10_ (0.05/*n* and 0.1/*n*), where *n* is the number of SNPs) were used to determine significant and suggestive associations between SNPs and phenotypes, respectively.

### Quantitative trait loci mapping

QTL analysis was performed with the R/qtl package ^[61]^ using the linkage map and marker data for Caméor × JI0281 RILs, as described by Ellis et al. ^[55]^. The composite interval mapping (CIM) method was used with a default window size of 10 cM and 3 background control markers. The significant LOD threshold at alpha = 0.05 for each protein band was determined after 1000 permutations. The QTL interval and proportion of phenotypic variance associated with the SNP with the highest LOD score were calculated using functions in the qtl package.

## DATA AVAILABILITY

The datasets used in the current study are available from the corresponding author upon reasonable request.

## AUTHOR CONTRIBUTIONS

J.B. and C.D. conceived the project and the research plans. A.O.W. performed the experiments, analysed the data, and drafted the manuscript. All authors reviewed the manuscript.

## ADDITIONAL INFORMATION

The authors declare that they have no conflicts of interest.

## ACKNOWLEDGEMENTS

We thank our colleagues at the John Innes Centre for their assistance during this work. We are grateful to Noel Ellis and Julie Hofer for providing seeds of the RIL population; Darryl Playford and Rebecca Lee for their help with field trials; and Carlo Martins and Gerhard Saalbach for running protein identification by LC-MS.

## FUNDING

This work was supported by the UK Department for Environment, Food, and Rural Affairs (grant number CH0111-CCN2; Pulse Crop Genetic Improvement Network). We also acknowledge the support from the UKRI BBSRC through the Institute Strategic Programme grant (BBS/E/J/000PR799).

## Notes

### Competing Interest Statement

The authors have declared no competing interest.

