## Supplemental figures 1-5 for "Identification of significant genome-wide associations and QTL underlying variation in seed protein composition in pea (*Pisum sativum* L.)"

**Supplementary figures**

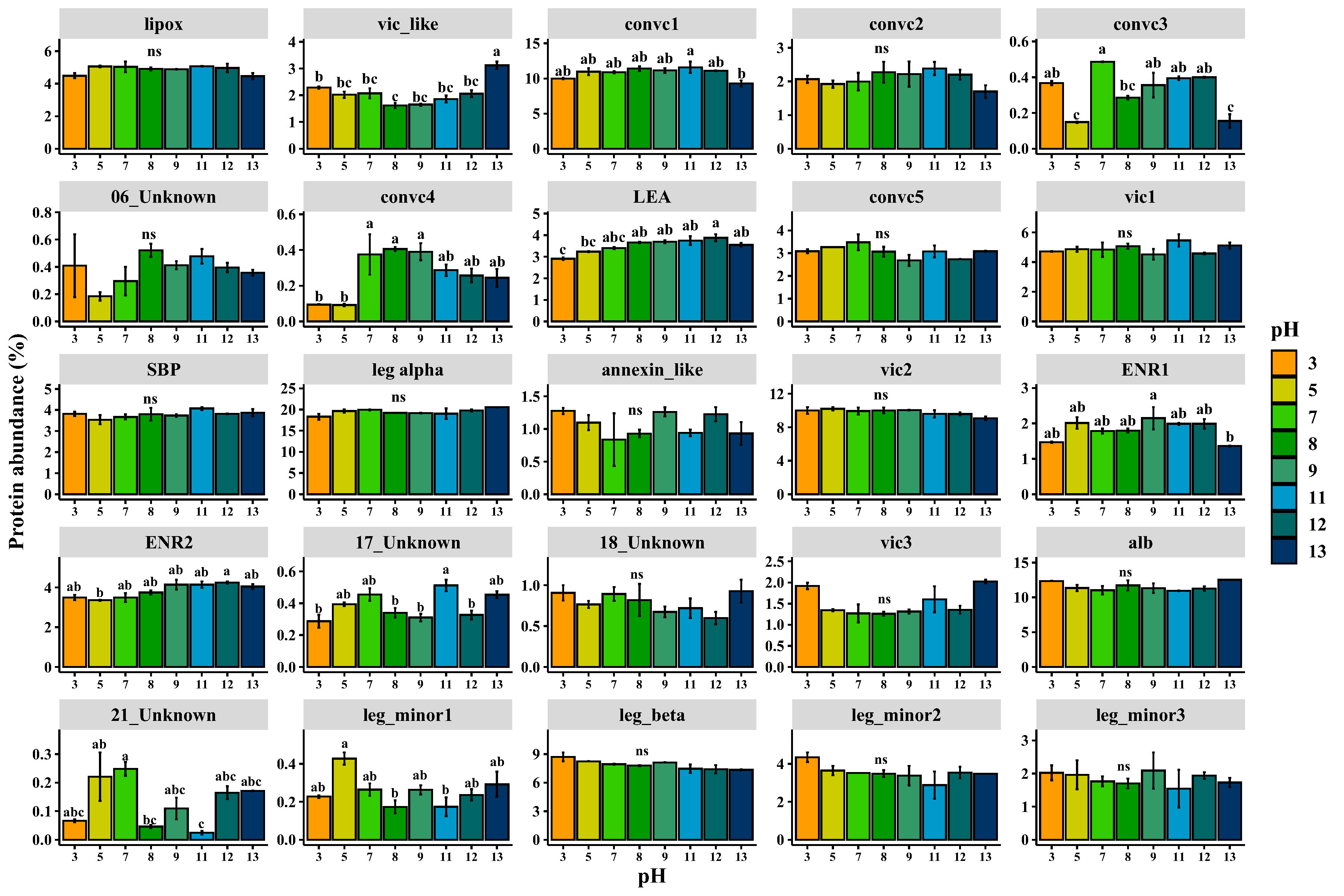

**Figure S 1**. Relative abundances of 25 protein bands quantified from pea seed proteins extracted with a buffer at eight pH levels. Total seed protein was extracted from a single pea accession, and the baseline buffer was 50 Tris/500 mM NaCl at pH 8, which was then adjusted with either HCl or NaOH. Tukey test was used to compare means, and different letters on the top of the bars indicate significant differences between treatments at ≤ 0.05 significance level. Means followed by the same letter are insignificant, while “ns” means no significant change across all the tested conditions. Error bars indicate the standard error of the mean.

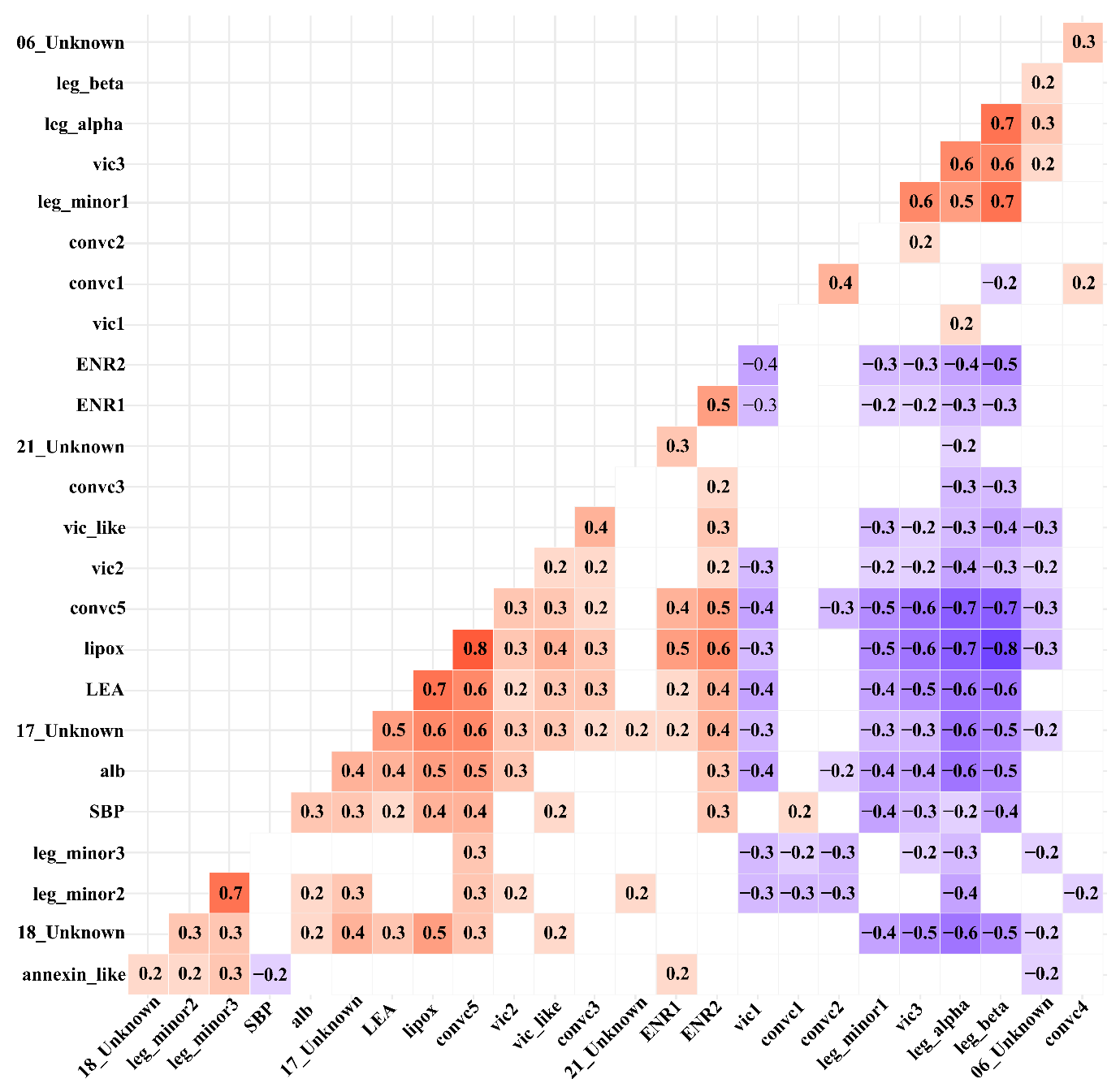

**Figure S 2.** Heatmap of Pearson correlation coefficient between seed proteins using 196 accessions grown in 2021 and 2022. The red and blue colours represent positive and negative correlations, respectively, whereas the intensity of the colours indicates the strength of the correlation. Only pair-wise correlations that were significant (p ≤ .05) are presented.

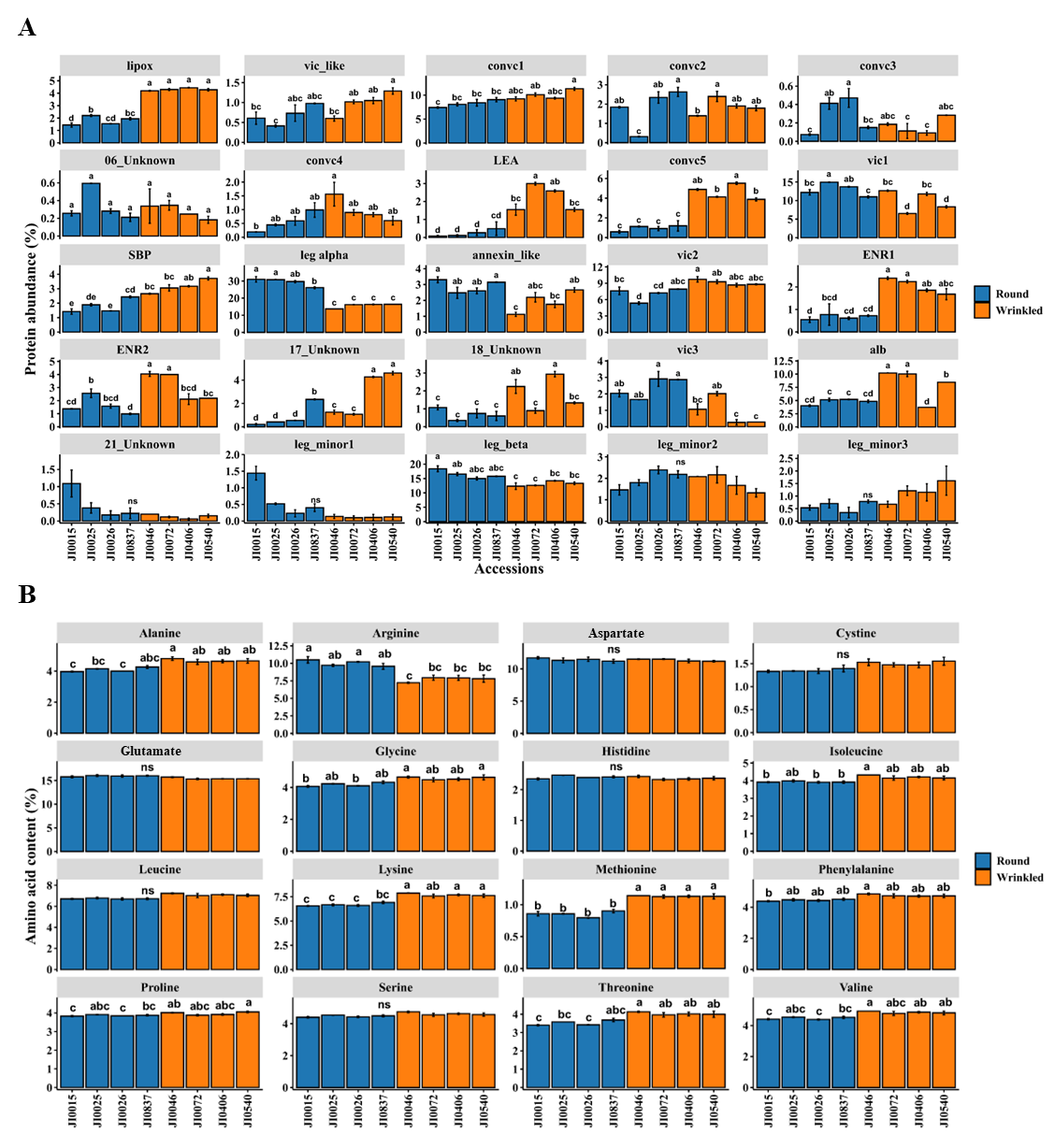

**Figure S 3.** (A) Eight pea accessions of round and wrinkled seeds with significant differences in the relative abundance of seed proteins. The blue and orange colours represent round and wrinkled peas, respectively. (B) Amino acid composition (% protein) of the selected eight accessions. The data are mean ± SE of two biological replicates from 2021 and 2022 harvests. Glutamic acid and glutamine were quantified as glutamate, and aspartic acid and asparagine as aspartate.

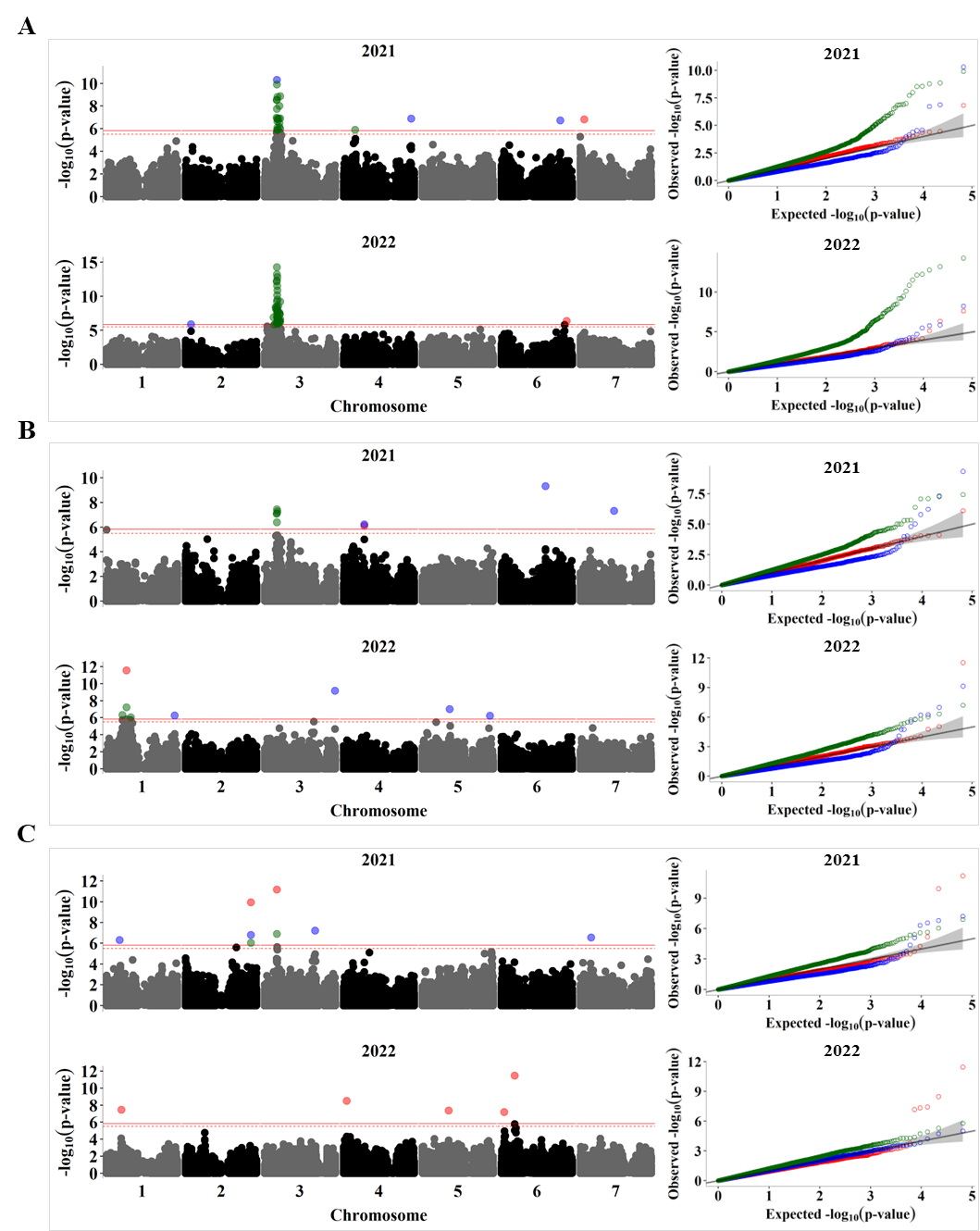

**Figure S 4.** GWAS analysis for the relative abundance of legumin polypeptides during the 2021 and 2022 seasons. Manhattan plots (left) and Quantile-Quantile (QQ) plots (right) for (A) leg_alpha, (B) Leg_beta, and (C) Leg_minor1. The solid horizontal line is the GWAS significance threshold corresponding to the false discovery rate (FDR) at p ≤ 0.05, while the dotted line is the threshold for suggestive association at p = 0.1. SNPs above the significance threshold in the Manhattan plots and the QQ plots are coloured by GWAS models, with red, blue, and green denoting results from BLINK, FarmCPU and SUPER, respectively.

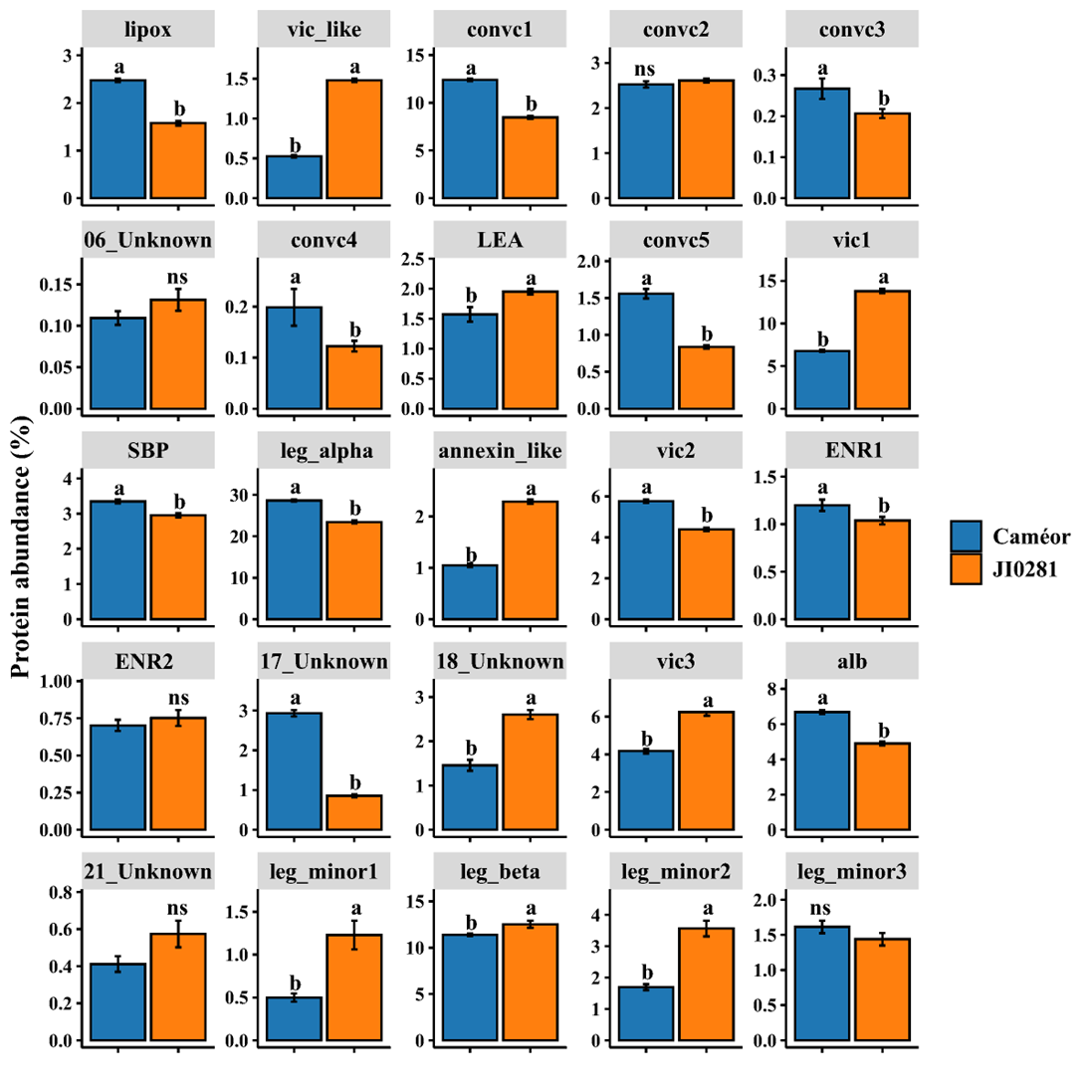

**Figure S 5.** Differences between the parental lines of the RIL population for the relative abundances of 25 protein bands. Different letters above the bars indicate statistically significant difference at p < 0.05 (t-test), while ns denotes no significant difference.
